# Meta-learning in head fixed mice navigating in virtual reality: A Behavioral Analysis

**DOI:** 10.1101/2023.05.01.538936

**Authors:** Xinyu Zhao, Rachel Gattoni, Andrea Kozlosky, Angela Jacobs, Colin Morrow, Sarah Lindo, Nelson Spruston

## Abstract

Animals can learn general task structures and use them to solve new problems with novel sensory specifics. This capacity of ‘learning to learn’, or meta-learning, is difficult to achieve in artificial systems, and the mechanisms by which it is achieved in animals are unknown. As a step toward enabling mechanistic studies, we developed a behavioral paradigm that demonstrates meta-learning in head-fixed mice. We trained mice to perform a two-alternative forced-choice task in virtual reality (VR), and successively changed the visual cues that signaled reward location. Mice showed increased learning speed in both cue generalization and serial reversal tasks. During reversal learning, behavior exhibited sharp transitions, with the transition occurring earlier in each successive reversal. Analysis of motor patterns revealed that animals utilized similar motor programs to execute the same actions in response to different cues but modified the motor programs during reversal learning. Our study demonstrates that mice can perform meta-learning tasks in VR, thus opening up opportunities for future mechanistic studies.

## Introduction

One of the biggest challenges for the nervous system is to quickly solve tasks in complex natural environments, typically containing a large number of sensory cues, in order to maximize the animal’s chances of survival. It has been well established that associative learning, including Pavlovian conditioning and operant conditioning, powerfully modify animal behavior to produce favorable outcomes. However, forming simple cue-action associations alone is inefficient because the entire process needs to be repeated each time an animal encounters novel sensory cues, even if the underlying task structure remains unaltered. Instead, it is beneficial for the brain to extract information about general rules and apply them to new problems with similar task structures, regardless of specific sensory features. For example, in addition to remembering specific landmark features, a mouse may learn that rocks, but not fallen leaves, are more stable spatial landmarks, knowledge that can potentially accelerate learning in new environments.

Classical psychological studies have shown that animals indeed have the ability to accelerate the learning process based on experience. One of the pioneering studies to demonstrate this phenomenon was done by Harry Harlow in the 1940s (Harlow, 1949). Harlow sequentially trained monkeys in an object discrimination task with 344 different pairs of objects, one of which was rewarded in each pair. The monkeys started with trial-and-error exploration but learned faster and faster with more experience. Furthermore, Harlow tested the reversal learning in monkeys by flipping the rewarded object, after the original association was learned. The reversal learning was also accelerated after monkeys accumulated more experience with different sensory objects. Harlow’s study convincingly demonstrated that monkeys have the capacity of ‘learning to learn’. Harlow called this ‘formation of learning sets’, which is now more commonly called ‘meta-learning’.

Beyond its importance in psychology of learning and memory, meta-learning also interests machine learning researchers, as it is a critical component of general-purpose artificial intelligence (AI) systems (Botvinick et al., 2019; Langdon et al., 2022; Neftci and Averbeck, 2019; Wang, 2021; Wang et al., 2018).

Theories on reinforcement learning (RL), combined with modern AI network techniques, have made great success in achieving superior-to-human performance on well-defined tasks (Mnih et al., 2015; Silver et al., 2016). However, it is still challenging for an AI system to generalize in multi-task environments that require continual learning. Understanding how the brain achieves meta-learning may provide inspirations for the development of novel AI systems.

Despite decades of psychology literature, neuroscience research in meta-learning remains scarce, which hinders exploration of the underlying cellular and circuit mechanisms. Recent progress in neuroscience technologies, especially novel genetics, recording, and imaging tools in mice, has tremendously advanced the mechanistic understanding of cognitive functions. However, most studies on learning in mice still focus on Pavlovian and operant conditioning, primarily due to the lack of well controlled behavioral paradigms for more complex forms of learning, such as meta-learning. However, recent studies on ‘schema learning’ in rodents (Caglayan et al., 2021; Farovik et al., 2015; McKenzie et al., 2014; Samborska et al., 2022; Tse et al., 2007; Zhou et al., 2021) indicate meta-learning is not restricted to primates. These studies were all performed in freely moving rodents, mostly rats. Although freely moving behaviors are preferrable for many reasons (e.g., to monitor natural movements), head-fixed behaviors offer unique advantages for large-scale two-photon imaging (Sofroniew et al., 2016) and in-vivo whole-cell recording (Harvey et al., 2009). Head-fixed experiments can also be integrated with virtual reality (VR) systems, which facilitate control and flexible manipulation of sensory cues. Visual cues, used in the VR system, are particularly helpful in studying meta-learning since it is relatively easy to generate a large set of cues, compared to olfactory or spatial cues used in previous schema learning studies. However, there is widely shared concern that both head-fixation and vision dependence pose huge challenges to mice, based on the assumptions that unnatural movement during head-fixation might interfere with learning and that mice might have poor visual acuity. Therefore, it remains unclear whether meta-learning can occur in head-fixed mice performing VR tasks.

In the current study, we trained mice to perform a two-alternative forced choice (2AFC) task in VR. Using a sequence of multiple visual cues, we tested the animal’s capacity of meta-learning in two types of tasks: generalization and serial reversal learning, both resembling Harlow’s original report in monkeys. In both cases, mice showed significantly increased learning speed with more experience. Strikingly, the animal’s behavior underwent sharp transitions during reversal learning, with the transition point shifted earlier during serial reversals. We also analyzed the animal’s motor patterns while solving problems with different cues but the same task structure. Our analysis revealed that animals implemented similar motor programs to execute the same actions in response to different cues, but they used modified motor programs during reversals. Together, our results provide compelling evidence of meta-learning in head-fixed mice in VR and reveal interesting observations that will inform physiological studies in the future, thus offering an opportunity to elucidate the neural mechanisms underlying meta-learning.

## Results

### Behavioral paradigms

Head-fixed mice were trained to run on a spherical treadmill, with three large monitors in front of the animal to render the virtual reality scene. We modified a Y-maze task that we previously designed to investigate the neural mechanism underlying visually guided spatial navigation (Zhao et al., 2022). Mice ran through a linear corridor with a pair of distinct visual cues, each displayed on one side of the walls (Fig. 1A). One of the cues was associated with reward. Mice were trained to turn into the rewarded arm at the branch point. The configuration of cues (i.e., which cue was on the left/right wall) was randomized trial-by-trial (See Methods for details).

**Figure 1.**
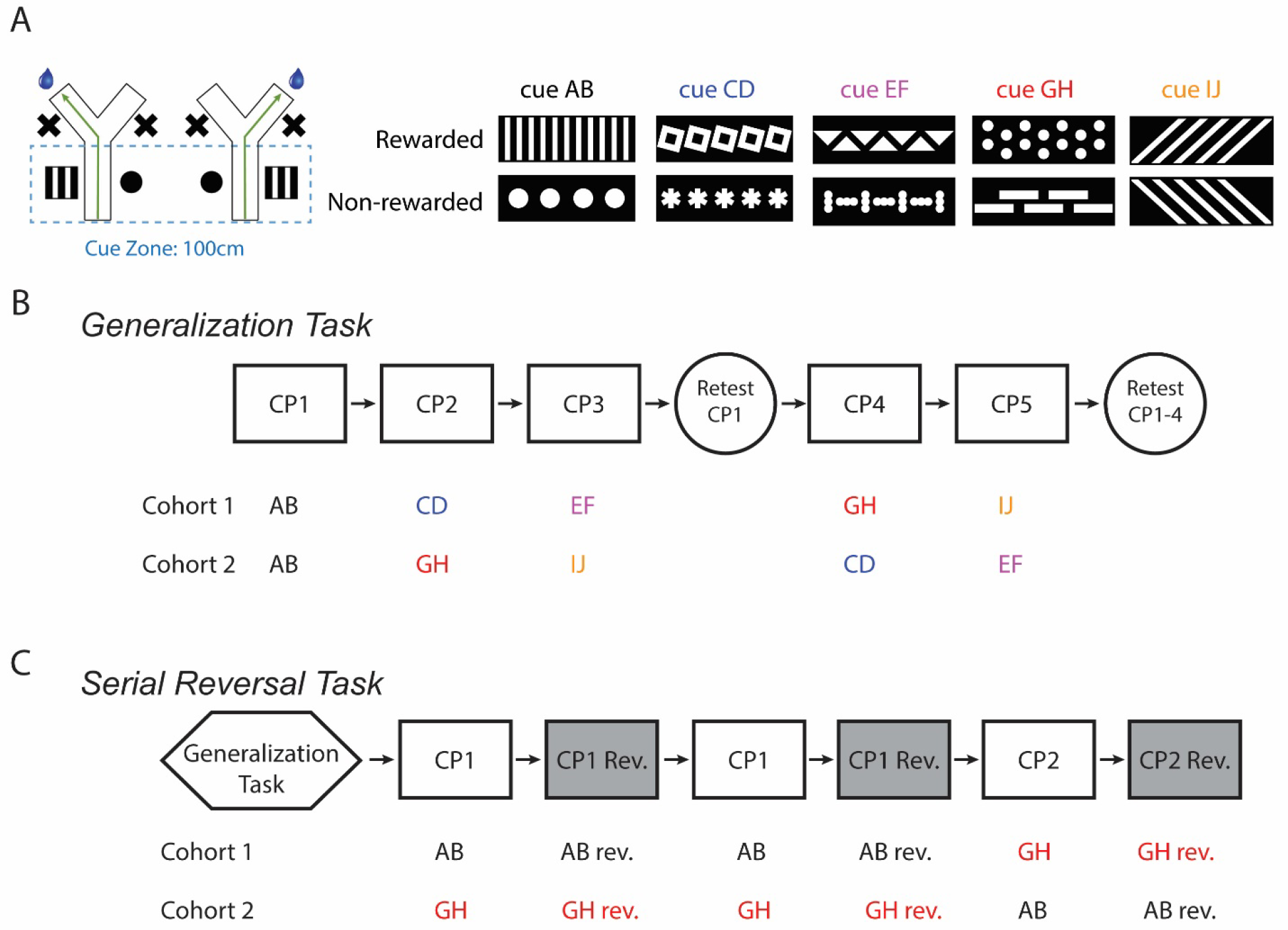
Design of behavioral paradigms **A**. Visually guided Y-maze task in virtual reality with 5 distinct cue pairs. One cue in each pair indicated the rewarded side. **B**. Generalization task. Two cohorts of mice were trained with counterbalanced cue- pair orders. The first cohort contained 6 mice while the second contained 5 mice. Each block represents multiple sessions. **C**. Serial reversal task. Two cohorts of mice (4 mice in each) were trained with cue pair AB or GH as the first reversal cue pair. Each block represents multiple sessions.

We designed two paradigms to test the mouse’s ability of meta-learning, a generalization task and a serial reversal learning task. In the generalization task, mice were sequentially trained with five different cue pairs (Fig. 1A). Each cue pair was used until the animal reached at least 80% success rate in one session. The first cue pair training started with the guided training mode as in our previous study (Zhao et al., 2022), in which a virtual wall was placed to block the entry point of the wrong arm, such that the animal would get stuck at the branch point until it switched to the correct direction. For later cue pairs, the animal was first trained in the regular version (both arms open) for 5-6 sessions. The training was switched to the guided mode if the animal did not reach the 80% criterion after 5-6 sessions. We introduced this guided mode because our preliminary experiments revealed that many mice could not learn the task within a reasonable amount of time when we just continued to train them in the regular Y-maze. The primary goal of this task was to characterize the animal’s learning speed at different stages of the whole training course. To minimize the confound that distinct learning speeds may result from different difficulties in discriminating various cue pairs, we counterbalanced the order of cue pairs in the training sequence (Fig. 1A). All animals started with AB as the first cue pair (CP1). One cohort of animals (6 mice) were trained with CD (CP2) and EF (CP3) first, followed by GH (CP4) and IJ (CP5), while the other cohort (5 mice) with GH and IJ before CD and EF (Fig. 1B). In addition to comparing learning speeds, we also tested whether animals could still remember previous cue pairs after learning new ones (i.e., avoid catastrophic forgetting). For this purpose, we implemented two retest blocks. One retest session, with the first cue pair (CP1), was held after the training of CP3. The last one or two sessions were broken up into a few short sessions to retest all cue pairs that animals had experienced.

All mice in the serial reversal task underwent the generalization task first. The rewarded and non-rewarded cues were flipped back and forth for two rounds with one cue pair (Fig. 1C). The reversal learning was then tested with another cue pair. To counterbalance potential different difficulties with different cue pairs, two cohorts of mice were trained. One cohort (4 mice) was trained with AB and GH as the first and second cue pairs, respectively, whereas the other cohort (4 mice) was trained in the reversed order (Fig. 1C). No guided training was performed in the serial reversal learning task.

### Mice generalized across cue pairs and avoided catastrophic forgetting

In the generalization task, it took a smaller number of sessions for animals to learn the fourth and fifth cue pairs (CP4 and CP5) than the second and third cue pairs (CP2 and CP3), in both example animals (Fig. 2A and 2B) and the population (for CP2-5: 9.4 ± 1.6 sessions, 10.7 ± 1.8 sessions, 3.4 ± 0.3 sessions, 3.9 ± 0.7 sessions, Fig. 2C). We did not compare CP1 with later cue pairs, since animals were required to adopt unique behavioral adjustments in CP1 sessions, such as adaptation to the head-fixation and learning to run and turn on the spherical treadmill. The analysis of session numbers might be complicated by the inclusion of guided training, which might not have the same impact on the animal compared to the regular training. To quantify the change of learning speed more rigorously, we focused on the animal’s performance with each cue pair when implemented in the early (as CP2 and CP3) versus late (as CP4 and CP5) phases of the training course. As expected, different cue pairs showed distinct levels of difficulty for mice to learn, with GH, IJ, CD, and EF being the easiest to the most difficult (as seen in success rates when trained in the early phase, Fig. 2D). Success rates were calculated in session 3 for CD and EF and session 2 for GH and IJ, as only 2 sessions were run with GH and IJ in some animals due to rapid learning. Quantifying success rates for all cue pairs in session 2 did not qualitatively change our conclusions (Supplementary Fig. 1). Pairwise comparisons, showed that the success rate was significantly higher when the same cue pair was placed in the late phase for all cue pairs except GH (65.6 ± 3.3% vs. 78.8 ± 2.9% for IJ, 60.5 ± 4.8% vs. 84.6 ± 5.5% for CD, 51.8 ± 2.4% vs. 77.2 ± 5.1% for EF, early vs. late, Fig. 2D). This trend was also observed with cue pair GH, despite that it barely fell below the significance threshold (71.2 ± 4.3% vs. 89.3 ± 4.7% for GH, *p*=0.0519), likely due to the high baseline in the early phase since GH was the easiest cue pair. ANOVA analysis revealed significant impact of both cues and training phases (early vs. late) on the success rate.

**Figure 2.**
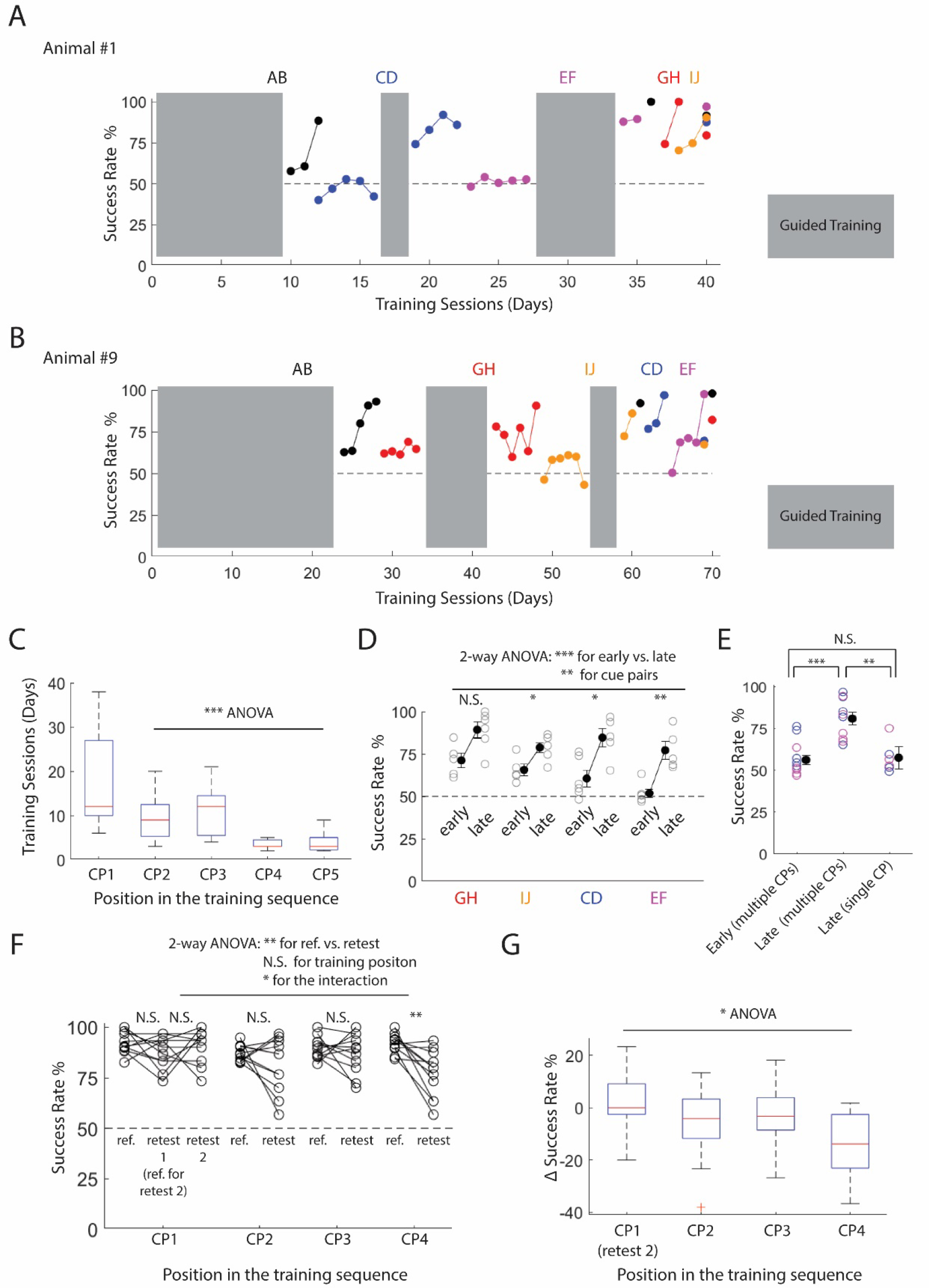
Generalization and avoidance of catastrophic forgetting **A**, **B**. Success rate in two example mice throughout the generalization task. Different cue pairs are color coded. Success rate was not calculated for the guided training sessions, as the animal was forced to turn into the correct arm. **C.** The number of training sessions required to reach 80% success rate. *p*=7.093e-5, CP2-CP5, ANOVA (n=11 mice). **D.** Success rate with each individual cue pair when placed in the early (as CP2 or CP3) vs. late (as CP4 or CP5) phases of the training. Data for GH and IJ were calculated from session 2, while CD and EF from session 3. *p*=0.0519, 0.0173, 0.0173, 0.0043, for GH, IJ, CD, and EF, early vs. late, Wilcoxon rank sum test (n=5 mice GH and IJ, early; n=6 mice for GH and IJ, late; n=6 mice CD and EF, early; n=5 mice for CD and EF, late). *p*=4.137e-8 for early vs. late, 0.0061 for cue pairs (2-way ANOVA). Black circles and error bars represent mean and S.E.M., respectively. **E.** Success rate with cue pair CD (blue) and EF (purple), in early and late phases in the multi-CP group (as in previous panels), and in the group with long-term single-CP training. p=6.819e-4, 0.0030, 0.4819 for Early (multi-CP) vs. Late (multi-CP), Late (single-CP) vs. Late (multi-CP), and Early (multi-CP) vs. Late (single-CP), Wilcoxon rank sum test (n=11, 11, 6 mice for early multi-CP, late multi-CP, and late single-CP). **F.** Comparisons of success rates between the reference and retest sessions. The reference session was the last session when the animal was trained with the same cue pair before the retest. CP1 had two retest sessions (Fig. 1). The first CP1 retest session was taken as the reference session for the second retest. *p*=0.2324, 0.4062, 0.2402, 0.5059, 0.0068, for CP1 retest 1, CP1 retest 2, CP2, CP3, and CP4, reference vs. retest, Wilcoxon signed rank test (n=11 mice). *p*=0.0077, 0.1124, 0,0116, for reference vs. retest, training positions, and the interaction of these two factors, 2-way ANOVA (comparing CP1 retest 2 and CP2-4 retest, n=11 mice). *p*=9.7656e-4 for all retest sessions compared with the chance level (50%), single sample Wilcoxon signed rank test (n=11 mice). **G.** Change of success rate from reference to retest. *p*=0.0247, 1-way ANOVA (n=11 mice).

The observed higher learning speed might be attributable to the animal’s experience with changing cue pairs. As a result, the animal may learn that this task contains flexible sensory features but always follows the same rule (one cue in a pair is rewarded). Alternatively, it could simply be the duration of training that sped up learning in the late phase. For example, the animal may improve its dexterity in maneuvering the spherical treadmill throughout training. To distinguish these two hypotheses, we trained a separate set of mice for a long period of time, but only with one cue pair (AB). 6 mice were trained with AB for 40 sessions, comparable to the mean duration of training by the end of CP3 in the previous set (21-65 sessions, 38.8 ± 4.9 sessions, mean±S.E.M., median 32 sessions). We then tested the animal’s performance with cue pairs CD (3 mice) or EF (3 mice). Interestingly, the success rate in this set was not enhanced, with success rates statistically equivalent to the early phase in the multi-CPs set (Fig. 2E). This result indicates that experiencing more than one sensory cue is critical for meta-learning.

To investigate the animal’s ability to avoid catastrophic forgetting, we measured success rates during retests for all cue pairs. Each one was significantly higher than the chance level (87.9 ± 2.4%, 90.9 ± 2.6%, 81.1 ± 4.1%, 87.1 ± 3.0%, 77.0 ± 3.6%, for CP1 retest 1, CP1 retest 2, and CP2-4, mean±S.E.M., Fig. 2F). The high performance in retest sessions was not due to quick re-learning of previous cue pairs within one session, as the rolling average of success rate did not significantly increase throughout the retest (Supplementary Fig. 2). Among all cue pairs, CP4 showed the largest reduction in success from reference to retest (Fig. 2G) and was the only cue pair that showed a significant decrease in success rate upon retest following learning of other cue pairs (Fig. 2F). This is surprising because CP4 had the shortest interval between the original training (reference session) and retest. One possibility is that animals were trained with CP4 for a much shorter time compared to CP1-3 (Fig. 2C); so the memory was less stable. Another possibility is that CP4 was learned after the animal already generalized the task structure, and thus fewer details of specific features were encoded (Robertson, 2018). These two explanations are related since memorizing details typically requires more time. Future studies are needed to distinguish these possibilities. Nonetheless, our results indicate that mice can generally avoid catastrophic forgetting, although there is moderate retrospective interference under some conditions.

### Mice exhibited increased learning rate through serial reversal sessions

As observed in both the example animal (Fig. 3A) and population (Fig. 3B), for CP1, the number of required training sessions to achieve at least an 80% success rate decreased in the second and third reversals compared to the first one (3.4 ± 0.6 sessions, 1.5 ± 0.2 sessions, 1.4 ± 0.2 sessions, for CP1 reversal, CP1 reversal back, and CP1 reversal 2, mean±S.E.M.). The two cohorts in the serial reversal task showed no significant difference and were thus pooled together in all analyses. We next calculated the 20-lap rolling average of success rate in all reversal sessions and defined the transition lap as the start lap of the first window in which the success rate increased to ≥80%. The cumulative number of laps to the transition lap showed significant reduction through the serial reversals (525 ± 87 laps, 177 ± 32 laps, 120 ± 12 laps, for CP1 reversal, CP1 reversal back, and CP1 reversal 2, mean±S.E.M., Fig. 3C). Strikingly, the increased learning rate was transferred to another cue pair, CP2, in terms of both the number of sessions and laps (1.25 ± 0.3 sessions, 167 ± 58 laps, Fig 3B and 3C). Together, these results demonstrate that mice exhibit meta-learning during serial reversal learning.

**Figure 3.**
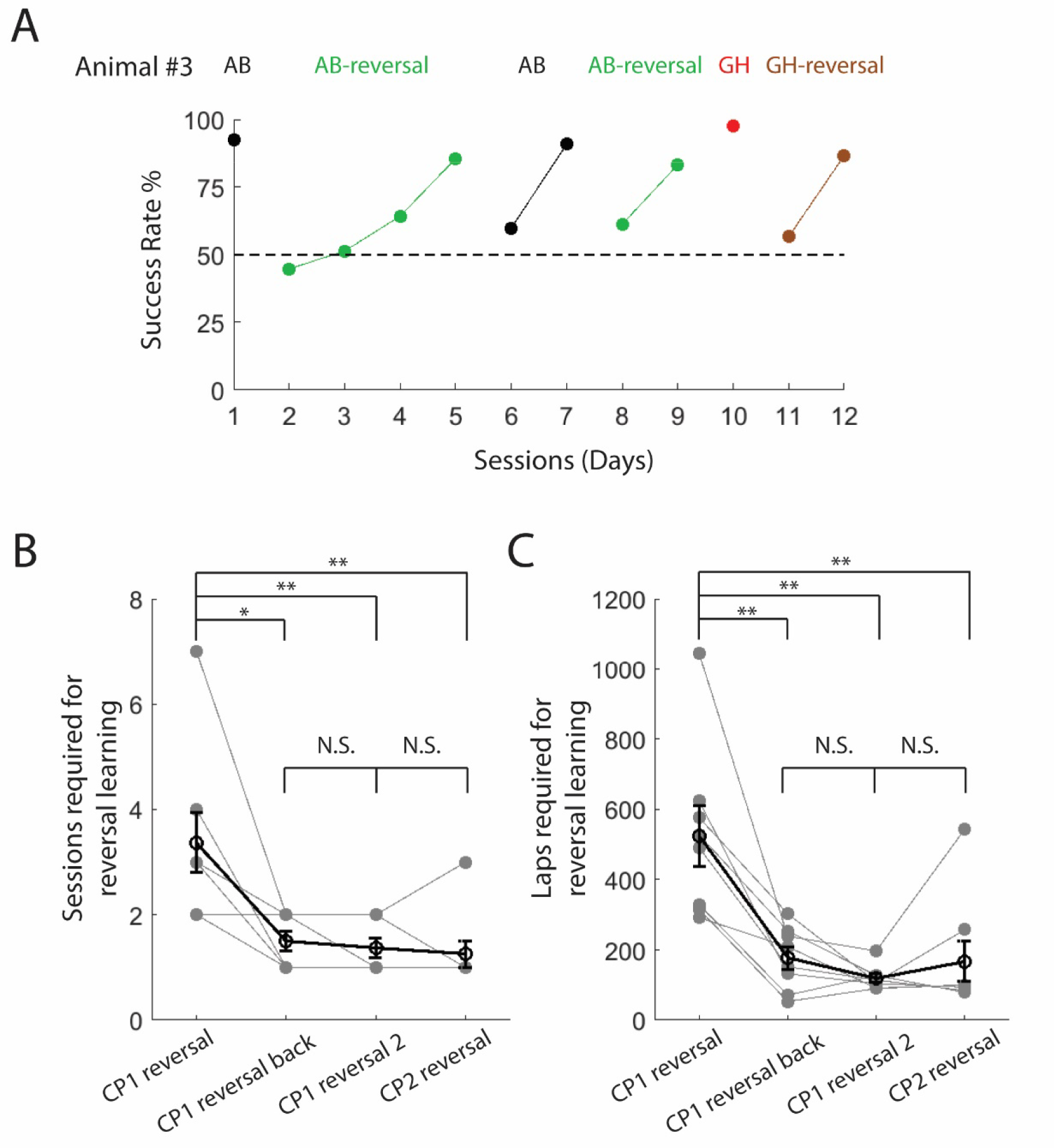
Meta-learning in serial reversal learning **A.** Success rate through serial reversals in one example mouse (AB as CP1 and GH as CP2). **B.** Number of sessions required for each reversal learning. *p*=0.0156, 0.0078, 0.0078, 1.0, 0.625, for CP1 reversal vs. CP1 reversal back, CP1 reversal vs. CP1 reversal 2, CP1 reversal vs. CP2 reversal, CP1 reversal back vs. CP1 reversal 2, CP1 reversal 2 vs. CP2 reversal, Wilcoxon signed rank test (n=8 mice). Black circles and error bars represent mean and S.E.M., respectively. **C.** Cumulative numbers of laps required for reversal learning (number of laps across multiple sessions were summed up). *p*=0.0078, 0.0078, 0.0078, 0.1484, 0.9922, for CP1 reversal vs. CP1 reversal back, CP1 reversal vs. CP1 reversal 2, CP1 reversal vs. CP2 reversal, CP1 reversal back vs. CP1 reversal 2, CP1 reversal 2 vs. CP2 reversal, Wilcoxon signed rank test (n=8 mice). Black circles and error bars represent mean and S.E.M., respectively.

We observed that mice showed discrete behavioral phases during reversal learning. As shown in the example animal (Fig. 4A), the animal’s success rate fell well below the chance level when the first reversal session started, because it still followed the original rule. However, the animal quickly changed its behavior after failing to get most rewards, by almost always turning to one direction, thus getting ∼50% of rewards (i.e., strongly biased turning). This may reflect the animal’s frustration and disengagement from the task. The biased phase could last for a long time, across multiple sessions (in this animal, the entire session 3 was in the biased phase). This behavior can be quantified with a large bias index, close to 100 (see Methods for details). At some point during session 4, however, the animal sharply transitioned out of the biased phase and began turning in both directions again, accompanied with a significant increase in success rate. Later reversals, exemplified by session 6 and 11 in this animal, showed similar transitions, although the biased phase lasted for a much shorter period (transitions occurred within one session).

**Figure 4.**
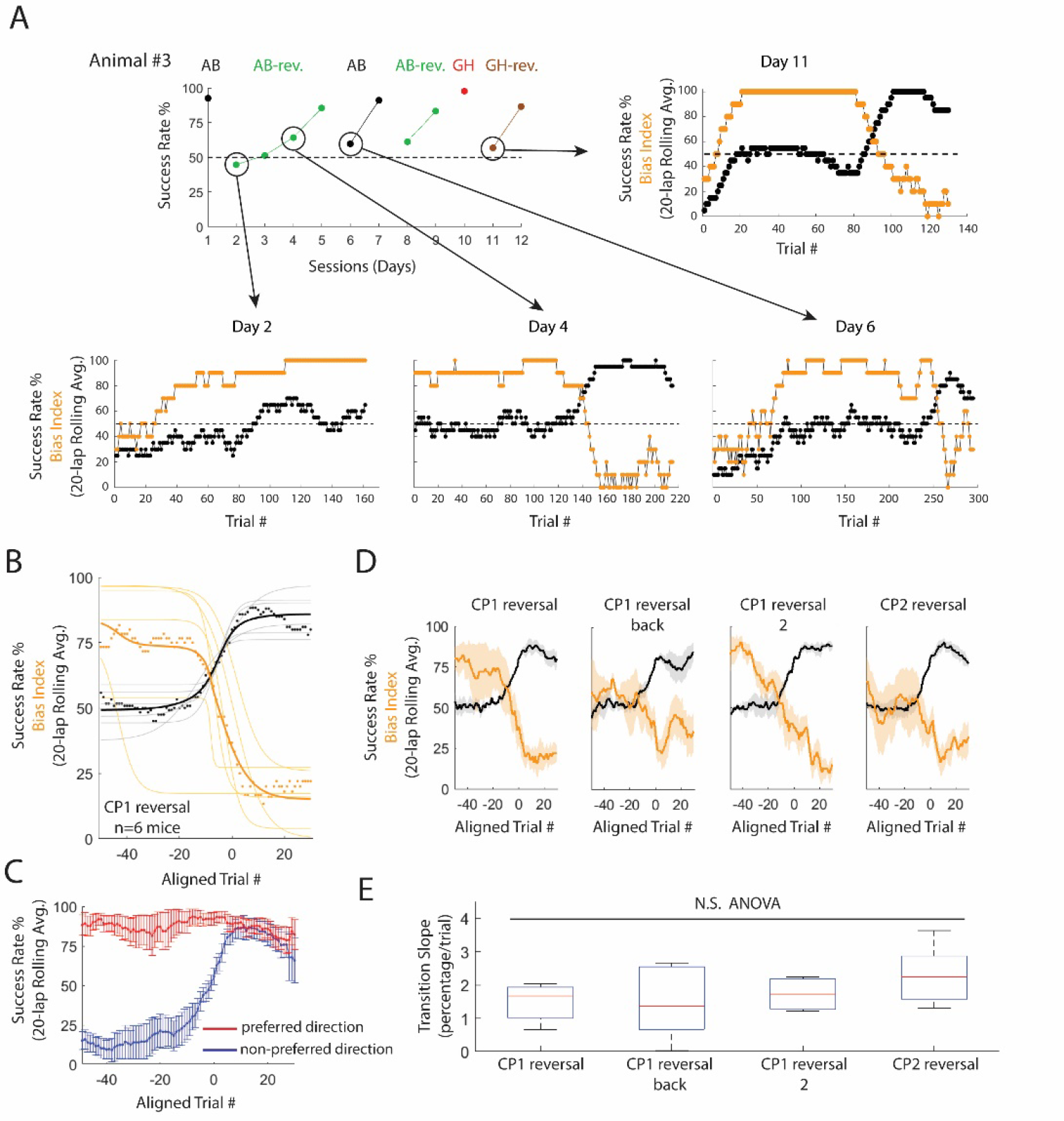
Sharp behavioral transitions during reversal learning **A.** 20-lap forward rolling averages of success rate (black) and bias index (yellow) in one example mouse during serial reversal learning. **B.** 20-lap forward rolling averages of success rate and bias index around the transition lap (lap 0) in all animals. Each light black and yellow curve represents fitted logistic function to an individual animal’s data. Dark black and yellow curves depict the average of all light curves. Black and yellow dots represent the mean of success rate and bias index, respectively, in all animals. 6 mice were included in this analysis. The other 2 mice had the transition lap too early in one session (<20 laps from the beginning) and were thus excluded since there were not enough pre-transition laps for the analysis. **C.** 20-lap forward rolling average of success rate for trials cued to the preferred (red) and non-preferred (blue) directions. Curves and error bars represent mean and S.E.M., respectively (n=6 mice). **D.** 20-lap forward rolling averages of success rate (black) and bias index (yellow) around the transition lap in 4 reversal sessions. Curves and shadings represent mean and S.E.M., respectively (n=6 mice). **E.** The slope of success rate change was calculated using the 10 laps before the transition lap (lap -10 to - 1, see Methods for details) in 4 reversal sessions. *p*=0.4018 (ANOVA, n=6 mice).

To quantify the transition from biased turning to successful turns in both directions, we aligned laps in all animals during the CP1 reversal sessions to the transition lap (defined as lap 0). The overall success rate sharply increased from approximately the chance level to ≥80%, while the bias index sharply decreased at the same time (Fig. 4B). In this process, the animal’s success rate in trials cued to the ‘preferred’ direction (the direction the animal turned during the biased phase) remained high, while the success rate in ‘non-preferred’ trials caught up (Fig. 4C). This transition happened within a very short period, ∼20 laps, compared to 525 laps on average required for the animal to complete the first CP1 reversal learning. Animals showed similar patterns of fast de-biasing and learning in all following reversal sessions (Fig. 4D). The increase in learning speed through serial reversals was not due to faster transition slopes (Fig. 4E), which was already very fast initially, but rather, the transition occurred earlier in the session.

The quantification of success rate and bias index both rely on the animal’s final turning decisions exiting the branch point. However, mice might carry out gradual modifications of motor patterns which appear to be ‘sudden’ in terms of success rate and bias index. For example, in the biased phase, the animal’s lateral running speed at the branch point in right-cued trials may start with a large left value and shift slowly to the right. At some point the lateral speed may cross 0 to become rightward, leading to an apparent ‘sudden’ switch although the underlying behavioral change is actually gradual. To test whether the change of motor pattern was also sharp, we focused on the ‘non-preferred’ trials since these were where the animal’s behavior changed dramatically. Our analysis revealed that the motor pattern also showed sharp transitions in ∼20 laps, similar to those quantified by success rate and bias index (Supplementary Fig. 3).

### Motor patterns in generalization and serial reversal learning tasks

We analyzed the animal’s motor patterns while solving generalization and reversal learning problems by quantifying the spatial profile of lateral running speed (‘roll’ of the spherical treadmill, see Methods for details).

In the generalization task, some animals maintained stable motor patterns throughout training, as shown in the example animal in Fig. 5A. The animal implemented similar motor patterns for all cues with only minor differences. In other animals, however, the motor pattern underwent significant changes, as shown in the example animal in Fig. 5B. This animal dramatically modified its motor pattern when training with cue pair GH. Interestingly, the newly adopted pattern was immediately applied to previous cue pairs on the retest day, without the requirement of extensive retraining with these cue pairs (Fig. 5B; see motor patterns for all animals in Supplementary Fig. 4). We quantified the correlation between motor patterns with the similarity index, using AB on the retest day as the reference (Fig. 5C; see Methods for details). Motor programs with most cue pairs on the retest day showed similar patterns (Fig. 5C). Statistical analysis revealed that the similarity comparing AB-initial and AB, with a long period of training in between, is significantly lower than the similarity across multiple cue pairs on the retest day (Fig. 5D). Finally, we tested the possibility that the similarity in motor patterns with different cue pairs was simply because mice operated the spherical treadmill in a highly stereotypic manner. This is unlikely the primary explanation of our observation, as the similarity across individual animals are significantly lower than across cues (Fig. 5E). Together, our results indicate that mice execute similar motor programs in solving the 2AFC task with different cue pairs and sometimes modify their programs during training.

**Figure 5.**
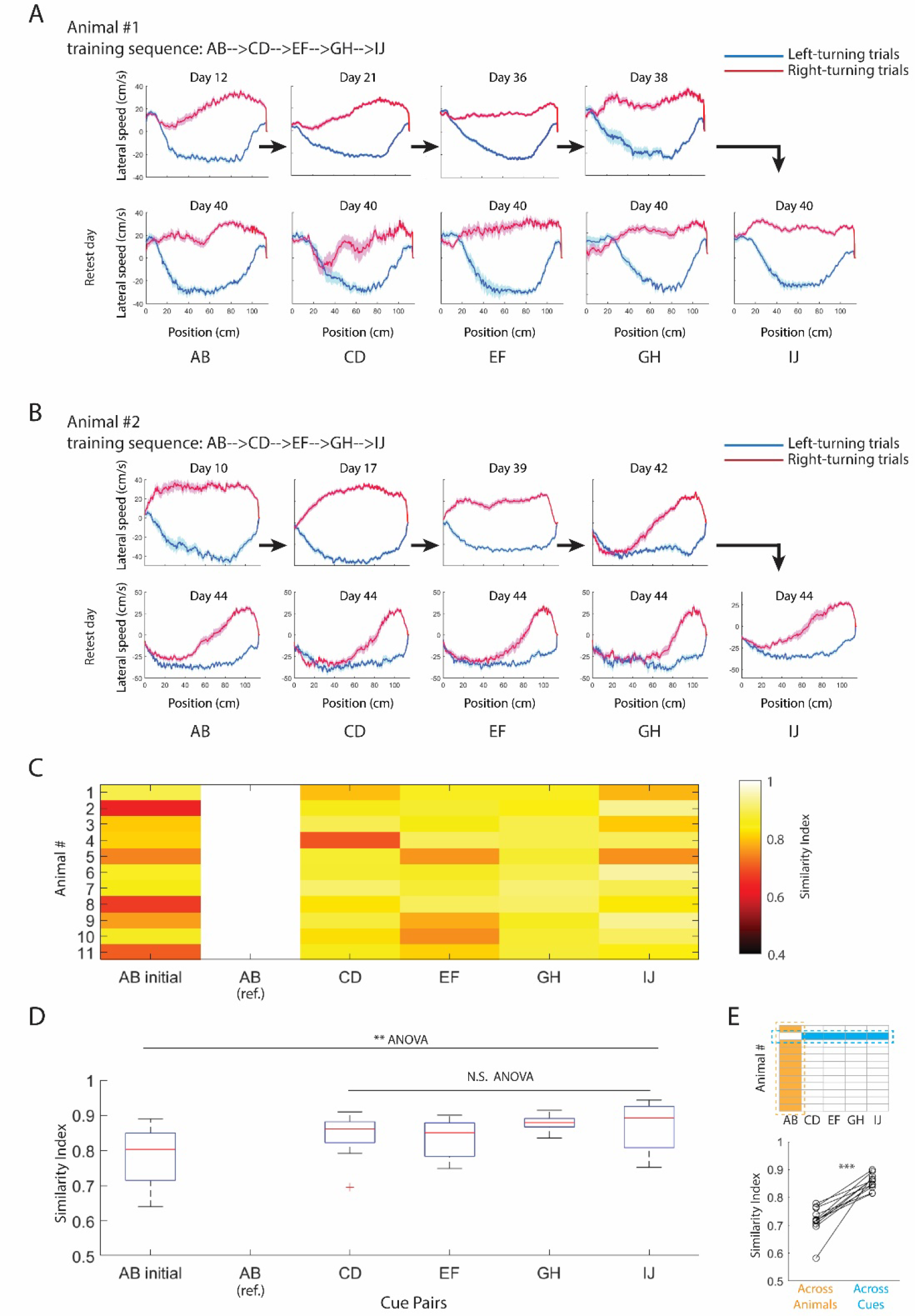
Motors patterns in the generalization task **A.** Lateral running speed through the trajectory in one example mouse (positive value, right turning; negative value, left turning). Red and blue represent right- and left-turning trials. Only correct trials were included in the analysis. Curves and shadings represent mean and S.E.M., respectively. Panels in each column are with the same cue pair. Day 12, 21, 36, 38, and 40 are chosen as they were the sessions in which the animal completed the training with cue pair AB, CD, EF, GH, and IJ, respectively. Day 40 was the final day of the training course in which all previously trained cue pairs were retested. **B.** The same representations as A in another example mouse. **C.** Similarity index of motor patterns in each animal, with AB in the same animal as the reference (see Methods for details). Motor patterns for AB, CD, EF, GH, and IJ were taken from the retest day, while from the last session of the initial AB training block (e.g., Day 12 and 10 in Animal #1 and #2, as in A and B) for AB initial. **D.** Statistics of motor pattern similarity index. *p*=0.0049 comparing all 5 groups, 0.2685 comparing the last 4 groups on the retest day (ANOVA, n=11 mice). **E.** Mean motor pattern similarity index across individual animals (with cue pair AB), and across multiple cue pairs on the retest day. Each pair of points in the figure represents one animal. The motor pattern with AB was used as the reference. Pairwise similarity indices were first calculated and then averaged in one column and row as the ‘Across Animals’ and ‘Across Cues’ data point, respectively (schematic cartoon). *p*=8.152e-5 Wilcoxon signed rank test (n=11 mice).

In the serial reversal learning task, most animals modified their motor patterns in reversal sessions, as shown in the example mice in Fig. 6A and 6B. Strikingly, the animal always implemented one pattern for ‘original sessions’ (i.e., sessions where the cue-reward association followed the rule as in the original training) and another for the ‘reversal sessions’ (i.e., sessions with cue-reward association reversed from the original rule), even with different cue pairs. This phenomenon was consistently observed in 6 out of 8 tested mice (Supplementary Fig. 5). To quantify this effect on the population level, we plotted the confusion matrix of motor patterns in 3 original sessions and 3 reversal sessions. It is clear that original and reversal sessions form two clusters (Fig. 6C). Motor patterns in CP1 original 2 and CP2 original were significantly more similar with CP1 original 1, compared to CP1 reversal 1, CP1 reversal 2, and CP2 reversal (Fig. 6D). Together, these results indicate that most animals created a new set of motor programs for reversal sessions, regardless of the specific cue.

**Figure 6.**
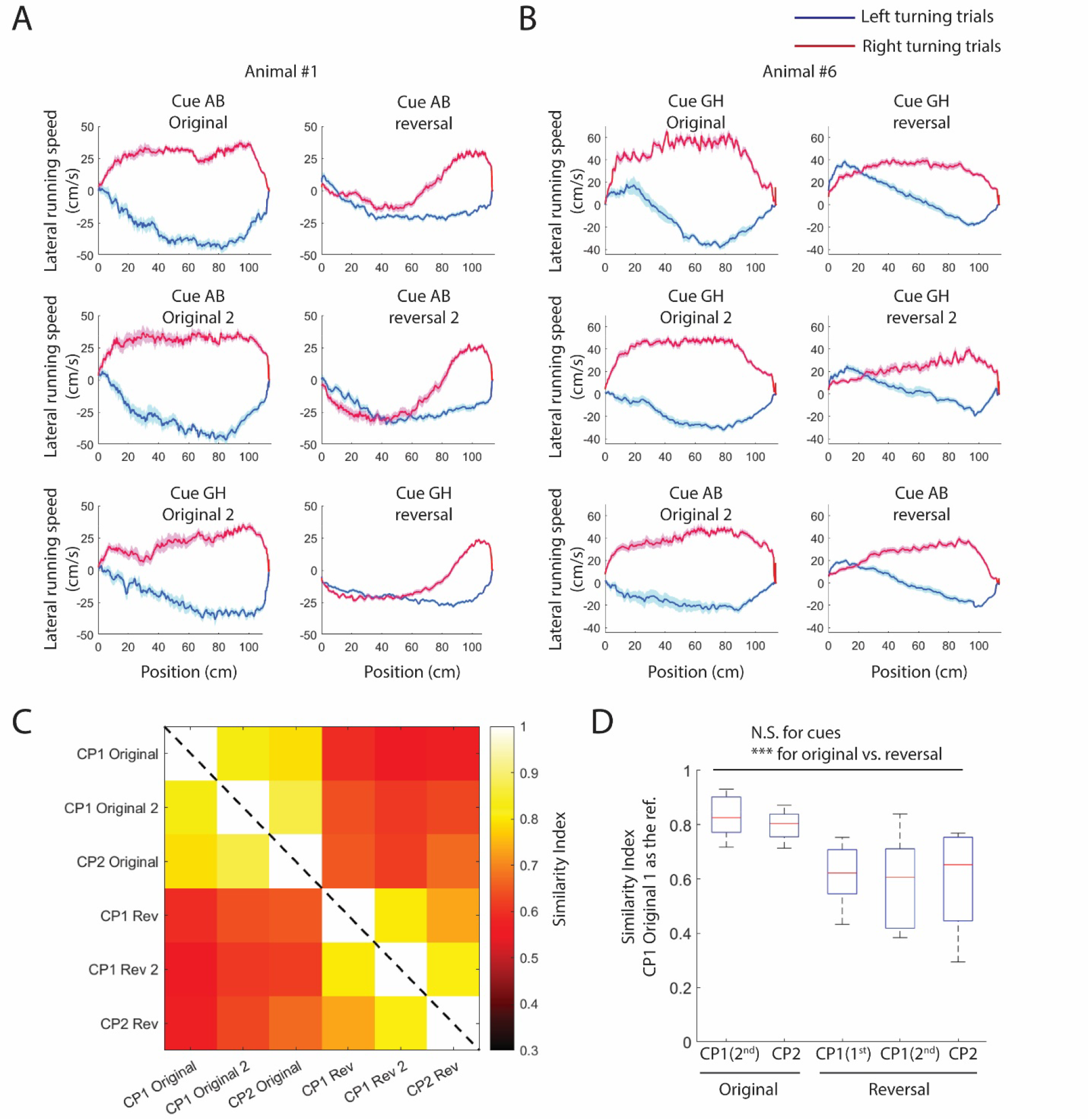
Motors patterns in the serial reversal learning task **A** and **B**. Lateral running speed through the trajectory in one example mouse (positive value, right turning; negative value, left turning). Red and blue represent right- and left-turning trials. Only correct trials were included in the analysis. Curves and shadings represent mean and S.E.M., respectively. **C**. Confusion matrix of motor patterns in 3 original sessions and 3 reversal sessions. **D**. Motor pattern similarity index in original and reversal sessions, with CP1 original 1 as the reference. *p*=8.127e-6, 0.6754 for original vs. reversal and CP1 vs. CP2, 2-way ANOVA (n=8 mice).

### Dorsal hippocampus is required for mice to perform non-delayed visually guided 2AFC in VR

Finally, we tested whether the dorsal hippocampus, a brain area well known for playing essential roles in memory and spatial navigation, is required for mice to perform the visually guided 2AFC task in VR. Our previous study revealed that the hippocampus is indispensable in a similar task (Zhao et al., 2022). However, the paradigm used in the current study has a critical difference: it has no delay zone. It has been well established that the hippocampus is required in contextual fear conditioning and trace conditioning, but not in cued conditioning without a delay period (Phillips and LeDoux, 1992). The hippocampus is not required in the spatial alternation task without delays either (Ainge et al., 2007). It is thus important to test whether the hippocampus is required in our current task, even in the absence of a delay period.

Bilateral silencing of the dorsal hippocampus was conducted by injecting fluorescently conjugated muscimol (Muscimol TMR-X) in both hemispheres. In 4 well trained mice, muscimol injections dramatically reduced the animal’s success rate to the chance level, while the effect was reversible 1 day later (Supplementary Fig. 6A). The animal’s locomotion, quantified by mean running speed (Supplementary Fig. 6B) and stop duration (Supplementary Fig. 6C), did not significantly differ under muscimol vs. control conditions. In 3 of these 4 mice, animals were still able to make turns in both directions in some epochs within the muscimol session, while the success rate stayed around 50% throughout the entire session (Supplementary Fig. 6D and 6E). Together, these results indicate that the disrupted performance in our memory task following hippocampal silencing is unlikely due to general interference of locomotion.

The most straightforward interpretation of the hippocampus dependence of our non-delayed paradigm is that it is a contextual memory task, i.e., the animal treats tracks with the same cue pair but opposite configurations (e.g. A-left/B-right vs. A-right/B-left) as two distinct environments. In a delayed Y-maze task, our previous study revealed global remapping in hippocampal pyramidal cells with different cue configurations (Zhao et al., 2022), consistent with the contextual switch. It is plausible that the delay is not required for animals to form concepts of different contexts. Therefore, our current non-delayed task may still be contextual. The behavioral paradigms, developed in this study, provide a great opportunity in the future to study whether the hippocampus, and associated brain areas, encode contexts based on the behavioral contingency (e.g., whether all left-turning trials with different cues are grouped as one context).

## Discussion

Meta-learning is an important feature of intelligence. It enables a flexible agent, animal or machine, to abstract task rules from experiences filled with sensory details and transfer them to solve new problems. Compared to human and non-human primates, neuroscience studies on meta-learning in rodents are limited. Previous studies on ‘schema learning’ and serial reversal learning have demonstrated meta-learning in rodents (Boulougouris et al., 2007; Bushnell and Stanton, 1991; Caglayan et al., 2021; McKenzie et al., 2014; Samborska et al., 2022; Zhou et al., 2021). In fact, schema learning is sometimes used interchangeably with meta-learning; however, meta-learning is a more general term for the ‘learning to learn’ phenomenon which does not necessarily require explicit representations of ‘schemas’ (relational structures) in certain neural circuits. Our current study is in general agreement with the schema learning literature and achieves three important advancements. (1) We demonstrate, for the first time, that mice can perform meta-learning in two visually guided VR paradigms (generalization and serial reversal learning). This opens opportunities to study meta-learning under head-fixed conditions with flexible online cue manipulations. (2) We designed cohorts with counterbalanced cue sequences, which is critical in providing rigorous evidence for meta-learning. Heterogenous baseline learning rate is a ubiquitous confound for measuring learning speed using sensory or spatial cues (e.g., rodents may respond differently to center vs. boundary positions in visual cues). In fact, we found that the impact of different cues on learning rate could be comparable to that of experience, emphasizing the importance of counterbalanced controls in studies on meta-learning. (3) We conducted detailed motor pattern analysis during meta-learning. Quantification of how an animal succeeds in getting reward in one trial provides more information that could facilitate interpretation of animal behavior and will also guide future mechanistic studies.

Simultaneous cognitive and motor processing is a general feature of diverse animal behaviors. One of the best examples is humans playing ball games, in which complicated strategic decisions and fine-scale movements must be made together. The major implication from our study is that the brain potentially encodes associative memories and motor programs (i.e., which action should be taken in response to a certain cue) in separate modules. This notion is supported by the following two observations: (1) Animals executed shared motor patterns for different cue pairs in both generalization and serial reversal tasks. 1. (2) In the generalization task, the associative memory of previously trained cues was preserved even after the motor pattern was modified during training with new cues. Notably, the novel motor patterns were immediately used in solving problems with old cues.

This observation has interesting implications for how sensory, memory, and motor systems interact during learning. Memory systems (e.g., hippocampus and prefrontal cortex) may form stable representations of cue groups with different behavioral contingencies (left-turning and right-turning), possibly arising from attractor dynamics. Sensory systems, likely visual cortices in our tasks, map individual visual cues to those established attractors in the memory system. The memory system then activates or biases the motor system (e.g., motor cortex, motor thalamus, and striatum) to drive behaviors. While learning new cues, the activated ensemble in the motor system sometimes shifted, while the attractors in the memory system were maintained unaltered, which explains why modified motor programs were immediately executed when animals saw previously trained cues (Fig. 7A). This hypothesis, if true, might be critical in explaining the increased learning speed during generalization since the formation of motor attractors in recurrent networks could be a slow process that does not need to be repeated with new cues. Furthermore, our motor pattern analysis in serial reversal learning showed that most animals implemented different motor patterns for original vs. reversed rules, while sharing the same set of patterns for different cues. Two hypotheses are consistent with this observation. First, the memory system may form a new set of representations for left- and right-turning trials, thus encoding original and reversed rules as two ‘hyper-contexts’ (i.e., higher level categories upon the left/right clusters) (Fig. 7B upper, hierarchical memory hypothesis). Alternatively, the contextual input about reversal may get integrated downstream to the memory circuit (Fig. 7B lower, contextualized motor hypothesis). Both models prevent original memories from being erased after reversal, which could explain the fast learning when the rule was flipped back. The key difference between these two hypotheses is whether representations in the memory circuit (e.g., hippocampus) shift during reversal. A recent study (Samborska et al., 2022), using a similar behavioral paradigm, reported minimal effects of reversal in CA1 representations, in favor of the contextualized motor hypothesis.

**Figure 7.**
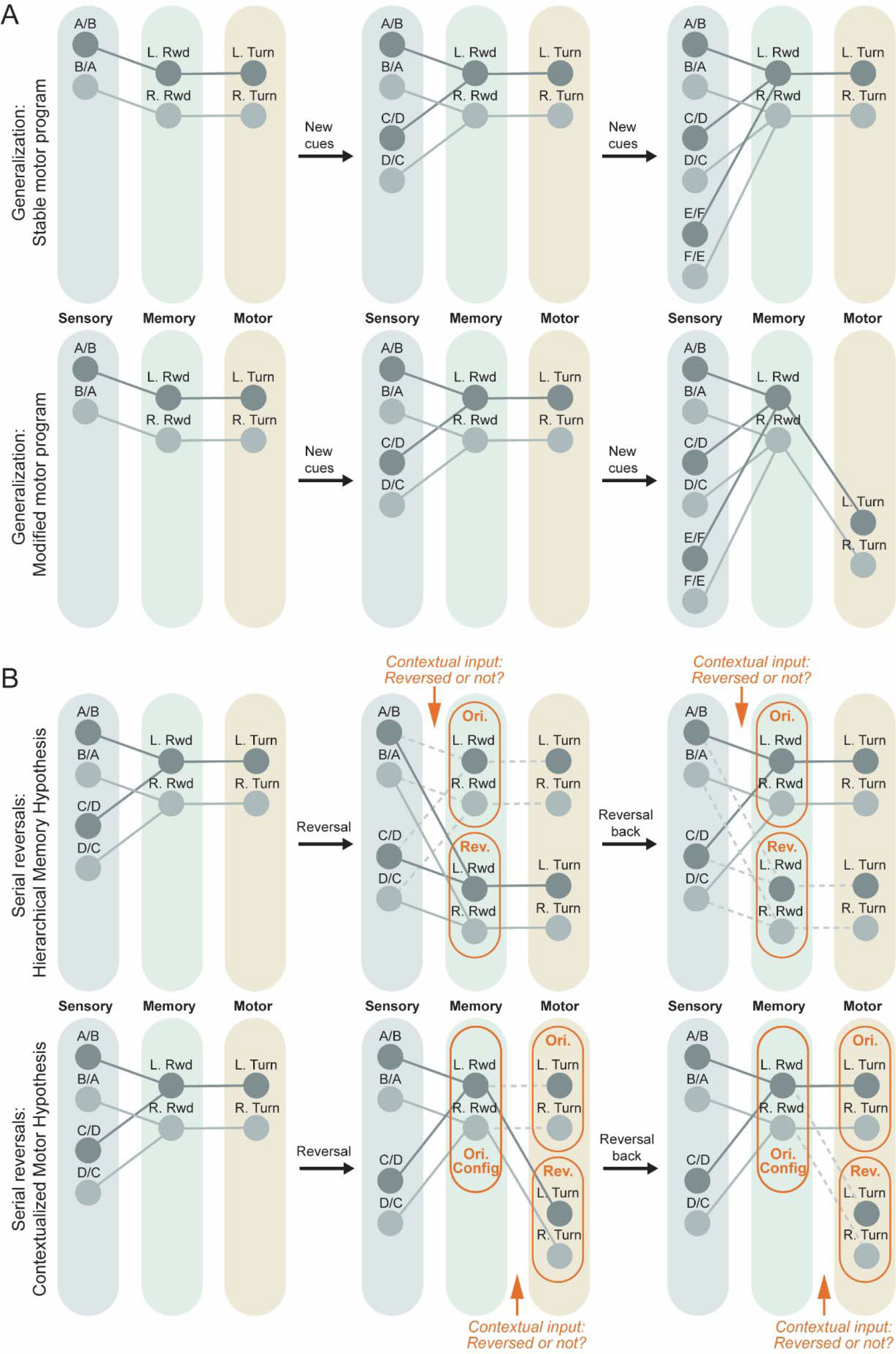
Working models for interactions between sensory (gray), memory (green), and motor (yellow) systems. **A.** The memory system forms representations of two reward locations, left reward (L. Rwd.) and right reward (R. Rwd.), during the learning of the first cue pair. These representations are preserved during subsequent cue pair learning. Two possible scenarios for the evolution of neural codes in the motor system during the generalization task are depicted. In the ’stable motor program’ scenario (upper panel), representations of the two motor programs, left turning (L. Turn) and right turning (R. Turn), remain unchanged throughout the entire learning process. In the ’modified motor program’ scenario (lower panel), representations of the two motor programs may be modified with new cue pairs. The modified motor programs are still mapped to the same reward representations in the memory system, allowing them to be immediately applied to all sensory cue pairs. **B.** Reversal learning does not eliminate the memory formed during the initial learning, but rather creates a new set of representations in either the memory system (upper panel, contextualized memory hypothesis) or motor system (lower panel, contextualized motor hypothesis). In the contextualized memory hypothesis (upper panel), a ‘contextual input’ indicating whether the current session is a ’reversal’ session is integrated into the memory system. Only one of the two memory ensembles, representing the original (Ori.) or reversal (Rev.) memory, is activated at a time, possibly through lateral inhibition. Distinct motor programs are mapped to the original and reversal memories. In the contextualized motor hypothesis (lower panel), the ’contextual input’ is incorporated into the motor system, selecting which motor program to activate. In this scenario, only one set of representations exists in the memory system. Dotted lines indicate suppressed pathways.

One striking phenomenon revealed in our study is the sharp transition of behavioral states during reversal learning. A recent study showed that freely moving mice can exhibit abruptly acquired ‘insight’ (Rosenberg et al., 2021). Our study demonstrates that sudden insight is not restricted to freely moving paradigms but can also be observed in head-fixed animals. In our reversal learning task, animals transitioned in and out of a biased behavioral state to eventually learn the rule reversals. This observation is consistent with a classic psychological study, which reported that rats, when presented with a reversal learning task, exhibited strongly biased turning followed by exploration behavior (described as “hypothesis” testing) prior to discovering the new task rule (Krechevsky, 1932). Rapid switch of behavioral preference during reversal has also been reported in non-human primates (Bartolo and Averbeck, 2020; Harlow, 1949; Hosokawa et al., 2018; Wilson and Gaffan, 2008), although the biased phase that we observed in mice was not typically observed. However, this discrepancy is likely explained by the fact that it only took a few laps for monkeys to learn the rule reversal. In fact, an early monkey study indeed reported hints of the biased phase in some slow learners (Cole, 1951), suggesting that our discovered phase transitions may be generalized to primates. Although it is reasonable to assume that the beginning of the biased phase reflects the animal’s frustration in not getting rewards in a presumably well-learned task, the biased phase may not be a continual disengagement period. For example, animals may exhibit some cognitive changes, such as enhanced attention to environmental cues or vicarious trial-and-error (Redish, 2016), that precede the animal’s transition out of the biased phase to above-chance task performance. Future physiological recordings will be helpful to identify potential changes in neural activity leading to this behavioral switch (Bartolo and Averbeck, 2020). Finally, it should be noted that the sudden shift in the animal’s behavior, or the output of the neural network, does not necessarily indicate that the underlying synaptic and cellular changes are abrupt themselves. Incremental weight changes often lead to stepwise reduction in the error, a well-known phenomenon in training artificial neural networks (Saxe et al., 2019). Both slow and fast (Bittner et al., 2017, 2015) forms of plasticity exist in the brain. However, the plasticity mechanism underlying sharp behavioral transitions remains to be investigated in the future.

It has been proposed that the hippocampus and frontal cortical areas (orbitofrontal cortex, OFC, and medial prefrontal cortex, mPFC) play complementary roles in meta-learning (Farovik et al., 2015; McKenzie et al., 2014; Samborska et al., 2022; Zhou et al., 2019), although the specific contributions of these brain area are still debated (see Zhou et al., 2019). It is hypothesized that rule representations in the frontal cortical areas, through slow learning, guide the quick formation of cue-action associations in the hippocampus. However, lacking manipulation and cellular physiology data, ideally throughout the training process, it is still unclear how the hippocampus and frontal cortical areas affect the firing pattern of each other at different stages during meta-learning. In the hippocampus, almost all previous studies on the integration of spatial and non-spatial inputs focus on the CA1 region, as it potentially integrates indirect inputs from frontal cortices, through the thalamus, and the spatial state representations generated within the hippocampus-entorhinal cortex (EC) complex (Frank et al., 2000; Ito et al., 2015; Samborska et al., 2022; Wood et al., 2000). Our previous studies, however, imply that the conjunctive code of spatial and non-spatial information occurred before CA1 (in CA3 or even more upstream) (Zhao et al., 2022, 2020). It is thus important for future studies to examine how CA3 and EC activity evolves during meta-learning and how inputs from frontal cortices affect them. The head-fixed VR meta-learning paradigms, developed in the current study, open opportunities for mechanistic studies to investigate these questions.

## Acknowledgements

We thank Drs. Mehrdad Jazayeri, Koichiro Kajikawa, Gabriela Michel, Kimberly Stachenfeld, Weinan Sun, and Yuhan Wang for their comments on the manuscript. We thank Julia Kuhl for her help with figure preparation. This work is supported by Howard Hughes Medical Institute and Tsinghua-Peking Center for Life Sciences.

## Author Contributions

X.Z. and N.S. conceived the study. X.Z., R.G., A.K., A.J., C.M., and S.L. conducted the behavioral experiments. X.Z. conducted the hippocampus silencing experiments. X.Z. analyzed the data. X.Z. and N.S. wrote the manuscript with input from other authors.

## Data Availability

Source data and analysis codes are deposited in a Figshare repository: DOI: 10.6084/m9.figshare.22725908

## Methods

### Animal Surgery

Adult (2-4 month old) male C57Bl/6 mice were used in all experiments (Charles River Laboratories). Headbar implant surgeries were performed as described before (Zhao et al., 2022). In short, mice were anesthetized with 1-2% isoflurane and mounted on stereotaxic (Kopf Instruments). A customized titanium headbar was implanted on the exposed skull with dental cement (Ortho-Jet, Lang Dental Manufacturing).

In experiments of bilateral hippocampal silencing, the coordinates for accessing dorsal hippocampus were marked during the headbar surgery (AP: -2.0, ML: ±1.7, from Bregma). Animals first underwent all behavioral training sessions in generalization and serial reversal tasks and were then probed with cue pair AB as the control session (counted as day 1). On day 2, the animal was anesthetized by isoflurane as in the headbar surgery, and small craniotomies on top of the dorsal hippocampus in both hemispheres were performed. The animal was then tested with cue pair AB following an 1h recovery in the home cage. On day 3, the animal was anesthetized and injected with 3.2mM BODIPY TMR-X muscimol conjugate solution (Invitrogen), as described before (Zhao et al., 2022). In each hemisphere, injections were made at 2 depths (1.25 mm and 1.65 from pia), 27.6 nL per depth, using a micro-injector (NanoJect). As on day 2, the animal was tested with cue pair AB following a 1h recovery. Procedures on both day 2 and day 3 took approximately 1h. On day 4, the animal was tested again with cue pair AB.

All experiments were performed according to protocols approved by the Janelia Research Campus Institutional Animal Care and Use Committee.

### Virtual Reality

Hardware and software used to control the virtual reality (VR) system has been reported in detail before (Cohen et al., 2017; Zhao et al., 2022, 2020).

A cued Y-maze task was adopted and modified from our previous study (Zhao et al., 2022). In the current study, the animal started at the beginning of a 100cm long track, with distinguishable visual cues displayed on both side walls. Cues were visible throughout the entire track. We removed the delay zone in the previous study to reduce the task difficulty. The animal then turned into one of the two arms (15cm) after the branch point. In each cue pair, there was one rewarded cue. The animal received a drop of 10% sucrose solution at the end of the arm if it made the correct choice. The left/right cue configuration was randomized trial-by-trial, except that a maximal number of three trials with the same configuration was allowed. In other words, if 3 left-cued trials occurred consecutively, the next trial would always be a right-cued one. We implemented this mechanism because, in preliminary experiments, we found that long repeating rows of the same configuration sometimes imposed disruptive effects on animals’ behaviors during training. In theory, this ‘three trial’ rule could affect the animal’s success rate in our tasks even if animals did not form cue-reward associations. However, our simulation revealed that the mean success rate is 57% (2000 sessions bootstrapping, data not shown), even if animals perfectly understand and act upon this ‘three trial’ rule (i.e., emit 100% correct responses in the next trial after 3 identical trials). Such perfect performance was never observed in any of our trained mice, and the 57% baseline does not qualitatively change any of our conclusions. Thus, we kept using 50% as the chance level for simplicity.

As in the previous study, the animal’s lateral location and head direction were locked at the center until it reached the branch point, from where the turning in the virtual environment was fully controlled by the animal’s locomotion. Rotations of the spherical treadmill on three axes (pitch, roll, and yaw) were monitored and recorded by optical flow sensors in our system. The pitch (forward rotation) was coupled with the animal’s forward/backward movement in VR. The roll/pitch ratio was used to determine the animal’s virtual turning.

### Behavioral Training

Animals were single housed after surgery. Five to eight days after surgery, animals were placed on water restriction, receiving 1.5 ml water per day. Body weight and health conditions were monitored every day throughout the entire experiment. Experiments were halted when the loss of body weight exceeded 30% or severe dehydration was observed.

Behavioral training started 1-2 weeks after the beginning of water restriction. Typically, one session (40-60min) was performed each day. Sessions were stopped when experimenters observed a reduction in the animal’s motivation, such as slow running, long pauses, or a lack of reward consumption. Animals typically ran 100-300 trials a day. In some animals, two sessions were conducted on transition days (from one cue pair to the next), including a short session with the old cue pair and then a longer one with the new cue pair. In the generalization task, the last 1-2 days were used to re-test all previous cue pairs (Fig. 1), with multiple short sessions on one day. On multi-session days, the animal remained on the set up while tasks were switched by changing environments in the VR system. Animals were not trained for 1-2 days each week due to experimenters’ work schedules, but water restriction was still implemented.

Once behavioral training started, each animal received 0.8-1.5ml water in total per day, depending on its motivation and tolerance to the water restriction. Water was supplemented if the animal did not receive enough in the task, as determined by weighing its overall health condition and its motivation to perform the task.

The first day of training only involved non-aversive handling and habituation to the head fixation, with the VR monitors off. From day 2 onward, animals were trained in the cued Y-maze in the VR. The speed gain was set high (2.5-3 folds as the real speed) initially to encourage running, and gradually reduced during training. The gain was fixed for each animal by the end of the first training block (cue pair AB). All animals started with the guided training, in which the wrong arm was blocked. The task was switched to the regular mode (both arms open), when the experimenter observed that the animal made frequent correct turns before hitting the blocking wall. The animal was then trained in the regular mode until it reached 80% success rate. Some animals showed poor success rates (<60%) in a few days, thus being switched back to the guided mode. Animals were moved to new cue pairs one day after reaching 80% success rate in one session.

In the generalization task, two blocks of retests were carried out, one after CP3 to retest CP1 and the other at the end of whole training course to retest all previously trained cue pairs (CP1-4, Fig. 1A). For the end-of-training retest block, 6 animals completed all re-tests in one day, whereas the remaining 5 completed the retests in 2 days, depending on the number of trials that they could run on one day. Retests were conducted following the reversed training order (CP4,3,2,1) in 6 mice while in forward order (CP1,2,3,4) for the other 5 mice. We did not observe significant difference between these groups and thus pooled them together.

Fourteen mice were trained in the generalization task. However, 3 of them dropped out due to health issues. Therefore, we included 11 mice in our current paper.

Eight out of the 11 mice were then utilized in the serial reversal task. All 8 of them completed the task. Before the serial reversal task started, we first re-trained these mice in cue pairs AB and GH (two cue pairs used in the reversal learning) for a few days to establish high baseline success rates. No guided training was used throughout the serial reversal task. Rule reversal (and reversal back) always started at the beginning of one session, not in the middle of a session. Animals were moved to the next phase when achieving 80% success rate.

### Quantifications and Statistics

Success rate was calculated as (number of correct trials)/ (number of total trials) X 100%.

To quantify the behavioral bias of the animal’s turning, we calculated a bias index defined as:

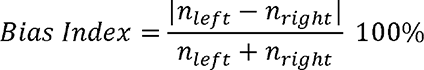

To analyze the change of success rate/bias index within one session, the rolling average of success rate/bias index was calculated in a forward window. In other words, the metric at trial n was defined as that metric in trials from n to (n+window length-1).

To analyze motor patterns, we focused on the lateral running speed (roll of the treadmill), since it determined the turning. The raw VR distance and speed data were first interpolated at 20k (original sampling rate was 60Hz, as the monitor’s refreshing rate). The speed was then smoothened by a Gaussian filter of 200ms S.D. The VR distance was binned with 0.1cm, and the mean speed was calculated within each spatial bin. Only correct trials were included, so that our motor analysis was not affected by differential success rates in different sessions.

To quantify the similarities between motor patterns (Fig. 5 and Fig. 6), we first normalized peaks of the lateral running speed in right-turning and left-turning trials as 0.5 and -0.5, respectively. The choice of 0.5 is because that the maximal difference between two points is 1. For right and left turning trials, we then calculated the mean squared distance between two curves:

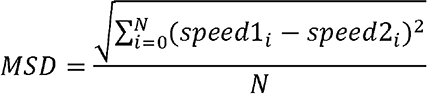

N is the number of spatial bins.

Since typically only one direction showed dramatic difference between sessions (Fig. 5B, Fig. 6A and 6B), we defined the motor pattern similarity index using the largest divergence from the two directions.

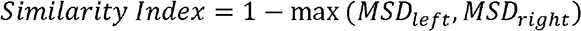

To analyze the evolution of motor pattern in the non-preferred direction during phase transition in reversal learning (Supplementary Fig. 3), we first calculated the correct trial and incorrect trial templates as the mean normalized lateral speeds of the first 5 correct trials and last 5 incorrect trials in our analysis window, respectively. We then calculated the normalized distance of each trial:

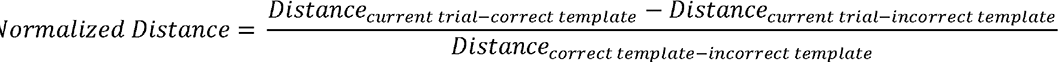

Fréchet distance was used in this calculation.

Since motor patterns before entering arms were important for the choice, we only used the motor pattern before the branch point (100cm) in this analysis.

To quantify phase transitions in reversal learning, we fitted our data with logistic function around the transition point:

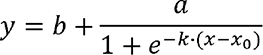

All statistical significance tests were conducted using non-parametric approaches. Wilcoxon signed-rank test and rank-sum test were used for comparing two groups in paired and unpaired manners, respectively. ANOVA was used for comparing multiple groups. In all figures, *, **, and *** indicate *p*<0.05, <0.01, and <0.001, respectively. Sample sizes were described in figure legends.

In all box-and-whisker plots, the upper/lower bounds of boxes and whiskers depicted 75/25 percentiles and maximum/minimum of the dataset, respectively. The central line indicated the median. Crosses were used to mark outliers, defined as data points outside the 1.5 times interquartile range. It should be noted that outliers were just labeled for the presentation purpose, not excluded from statistical analysis.

In all curve plots, the curve and error bar/shade depicted mean and S.E.M., respectively.

All quantifications and statistical analyses were done with customized MATLAB codes (version 2022a, Mathworks).

## Supplementary Figures

**Supplementary Figure 1.**
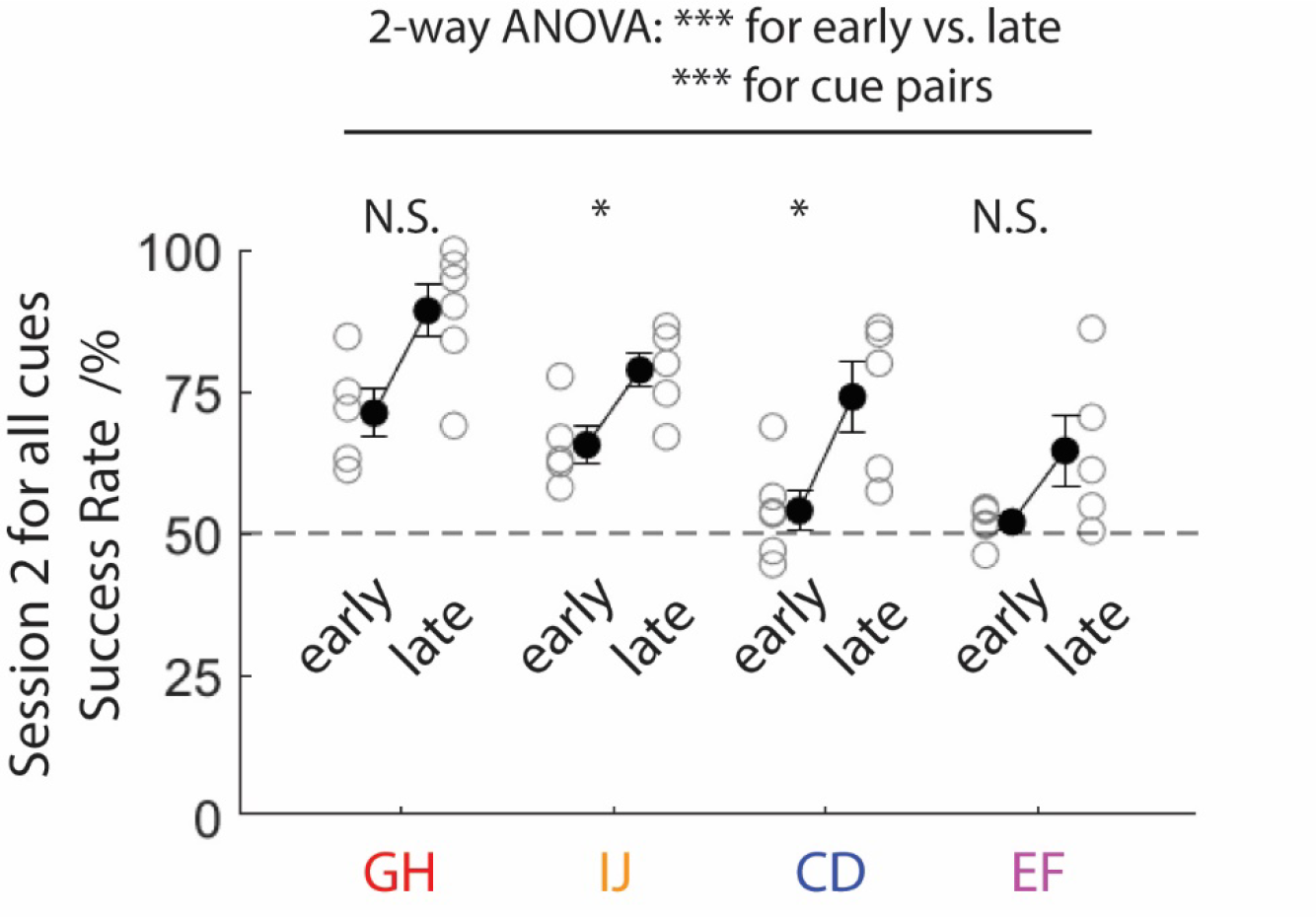
Performance on session 2 in early and late phases during the training Success rates on session 2 with multiple cue pairs, when placed in the early (as CP2 or CP3) or late (as CP4 or CP5) phase. The same as Fig. 2D except that performance data on session 2, instead of session 3, were used for cue pair CD and EF. *p*=0.0519, 0.0173, 0.0173, 0.0823 in pairwise comparisons for GH, IJ, CD, and EF, respectively (Wilcoxon rank sum test). *p*=2.431e-6 for early vs. late phases, and 2.324e-5 for cues in 2-way ANOVA analysis.

**Supplementary Figure 2.**
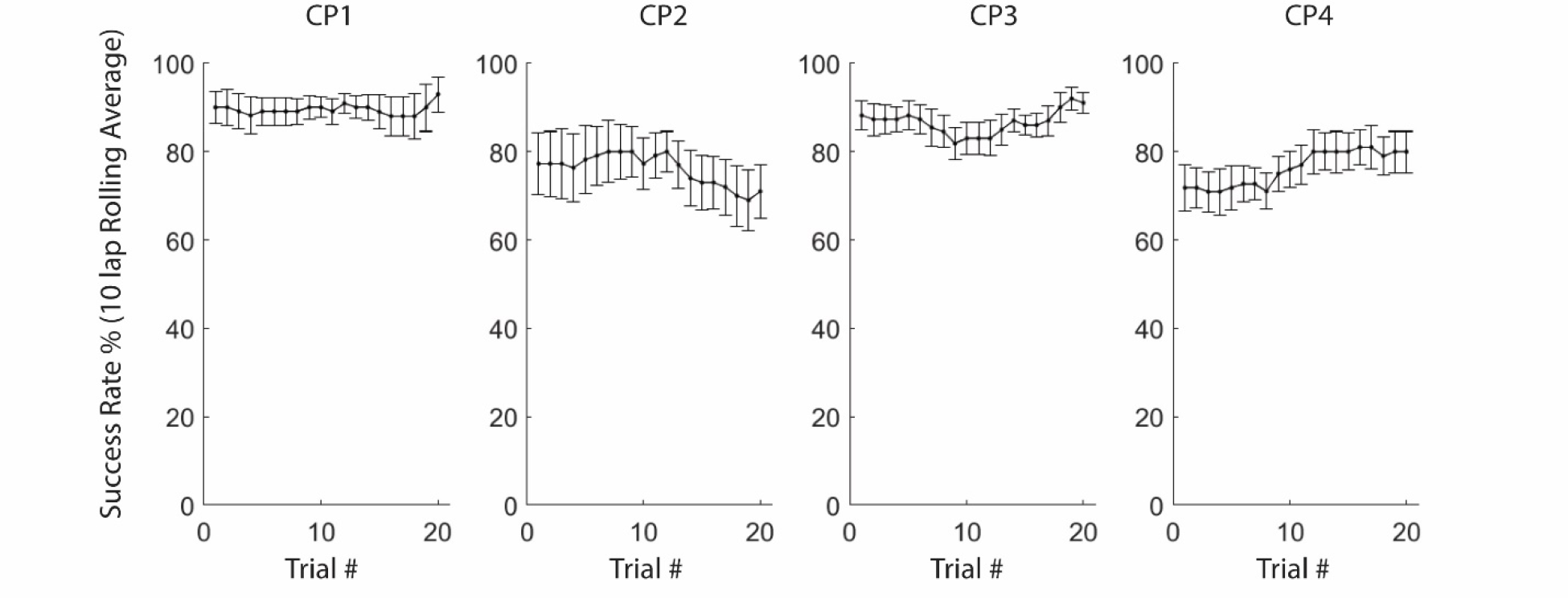
Success rate during retest sessions Success rates (forward 10-lap rolling average) during the retest at the end of the training course. X-axis presents the start trial number in each calculation window. Mean and S.E.M. were plotted (n=11 mice).

**Supplementary Figure 3.**
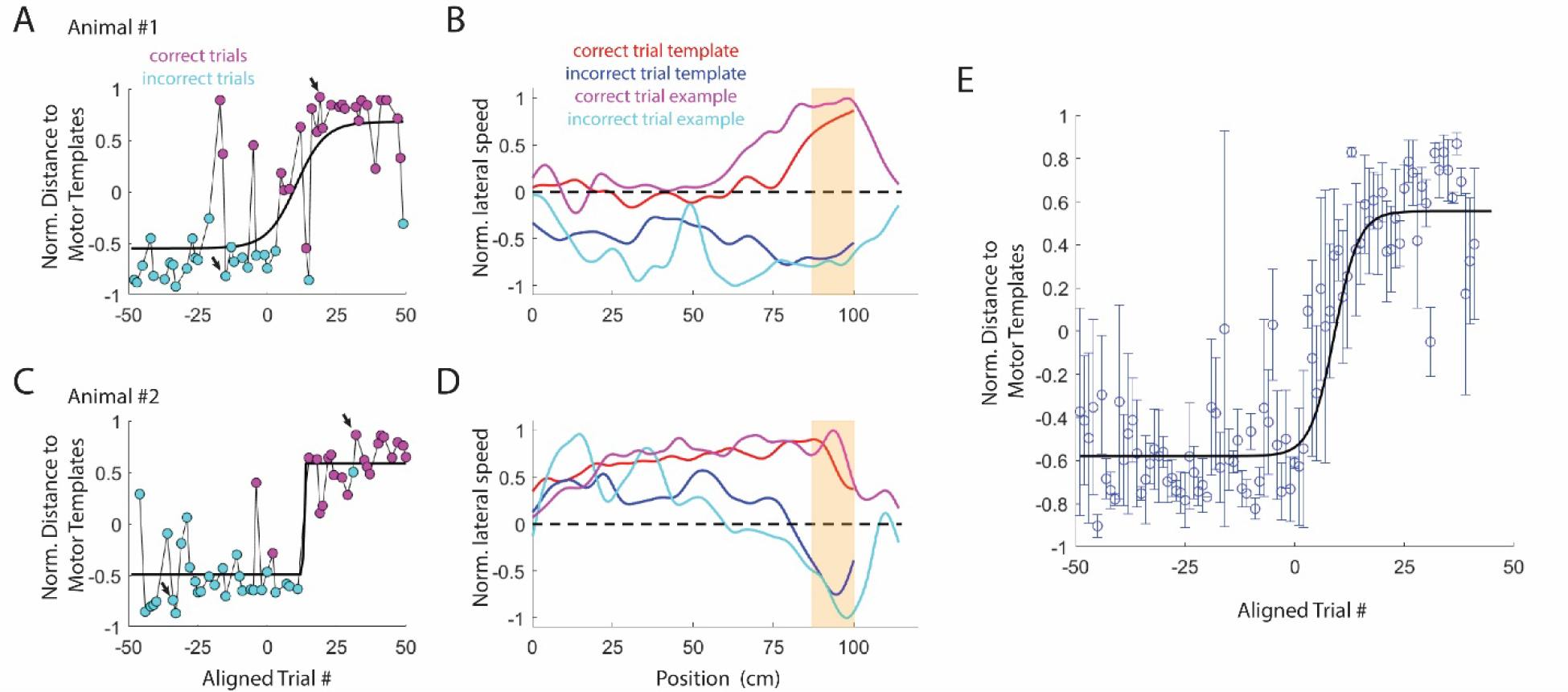
Transition of the motor pattern during reversal learning **A.** Normalized distance to motor templates in one example animal (-1 indicates the incorrect trial template while 1 indicates the correct trial template. See Methods for details). Magenta and cyan colors represent correct and incorrect trials, respectively. Lap numbers were aligned with the transition lap (the start lap of the first 20-lap window with success rate ≥80%). Black curves: fitted logistic functions. This analysis was focused on the non-preferred trial type, in which the cued direction was not taken by the animal during the biased mode. **B.** Templates of normalized lateral running speed for the correct (red) and incorrect (blue) trials. Normalized lateral running speeds for two example trials (arrows in A) were overlayed. Yellow shading: branch point of the Y-maze. **C** and **D**. The same as A and B in another example animal. **E**. Normalized distance to motor templates in all animals (mean±S.E.M., n=6 mice). Black curve: fitted logistic function.

**Supplementary Figure 4.**
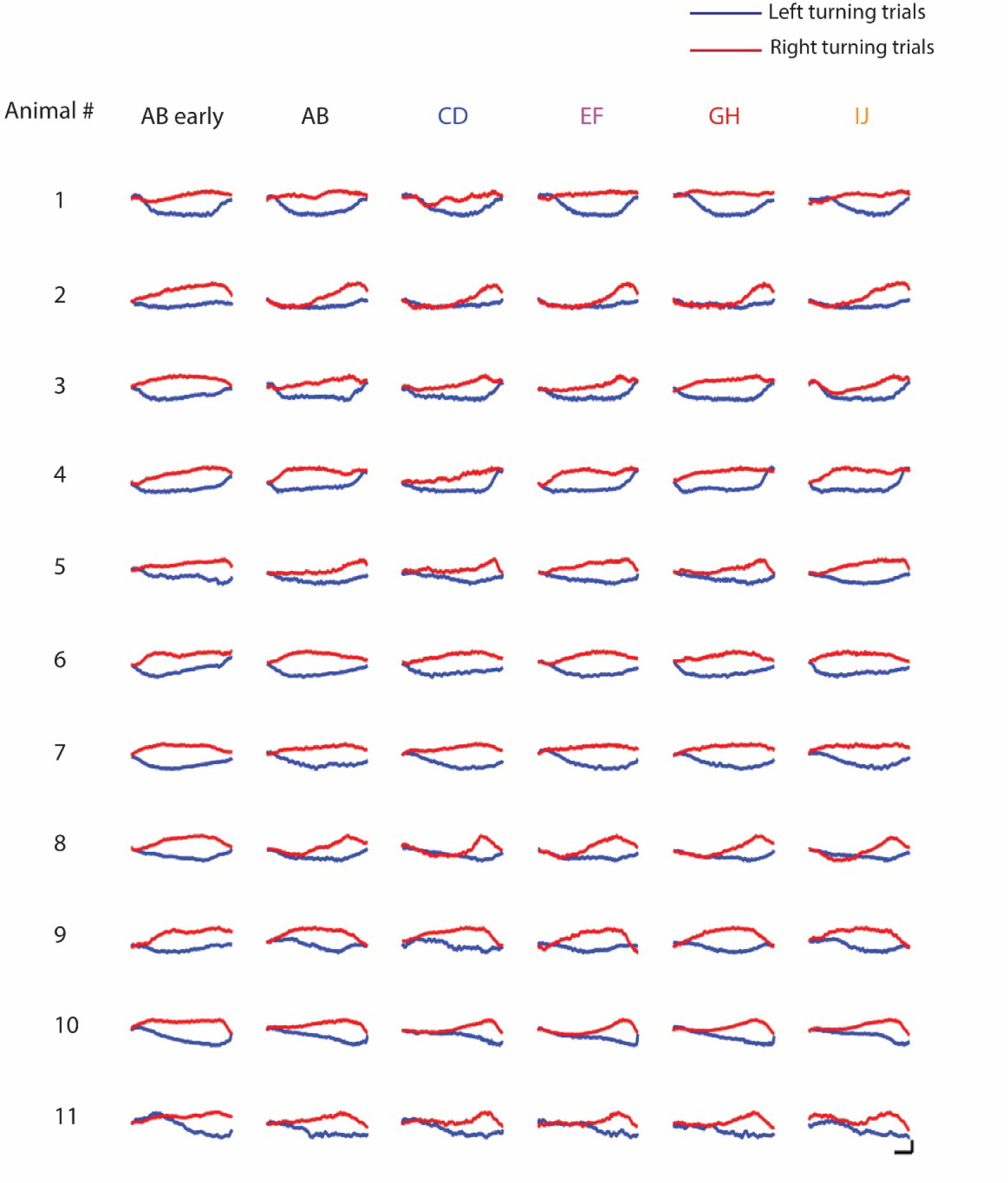
Motor patterns when tested with different cue pairs Position profiles of normalized lateral running (scale bars at the right bottom corner: 0.25 and 20cm). Blue and red represent left- and right-turning trials, respectively. The first column represents the session when the animal first learned to perform the task with cue pair AB in the initial segment of the training. The rest five columns represent the last one or two sessions when all cue pairs were retested.

**Supplementary Figure 5.**
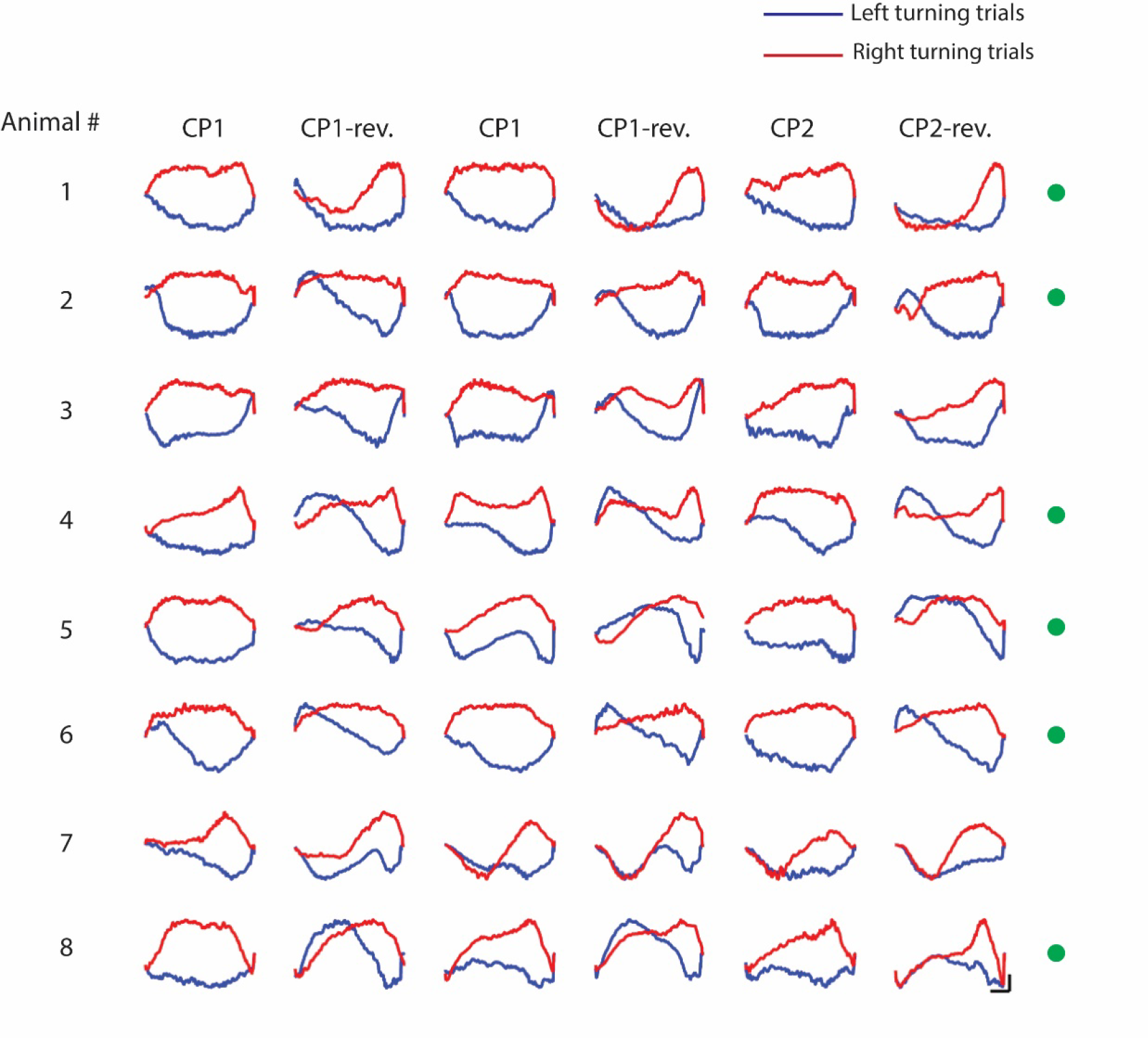
Motor patterns during serial reversal learning Position profiles of normalized lateral running (scale bars at the right bottom corner: 0.25 and 20cm). Blue and red represent left- and right-turning trials, respectively. Greens dots indicate animals where two distinct patterns for the original and reversal rules were clearly observed.

**Supplementary Figure 6.**
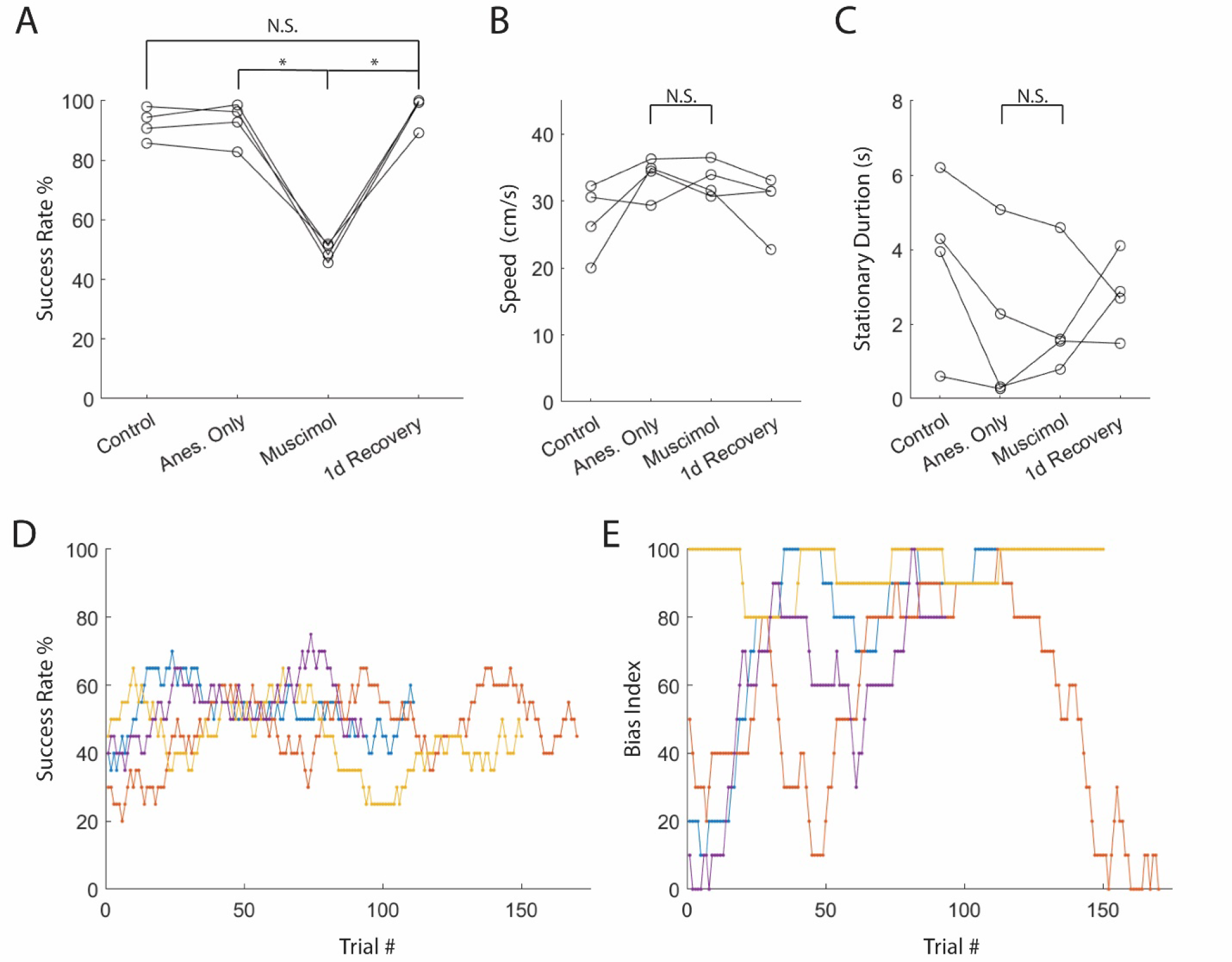
Silencing dorsal hippocampus disrupted the mouse’s performance on the visually guided 2AFC without delay zone. **A.** Success rates under different conditions, including control, 1h-recovery following 1h-anesthesia (Anes.-only), 1h-recovery following bilateral muscimol injections into the dorsal hippocampus (Muscimol), and 1d-recovery following the muscimol injections. *p*=0.0286, 0.0286, 0.1714 for Anes.- only vs. Muscimol, Muscimol vs. 1d-Recovery, and Control vs. 1d-Recovery, respectively (Wilcoxon rank sum test, n=4 mice). All mice were tested with cue pair AB. **B.** Locomotion speeds under different conditions. Mean speeds were calculated only from running periods, defined as sections where the animal’s speed was higher than 2cm/s. *p*=0.1143, Anes.-only vs. Muscimol (Wilcoxon rank sum test, n=4 mice). **C.** Durations of stationary periods, defined as sections where the animal’s speed was lower than 2cm/s. *p*=0.4857, Anes.-only vs. Muscimol (Wilcoxon rank sum test, n=4 mice). **D.** Success rate (20-lap forward rolling average) following muscimol injections. Different colors represent 4 different mice. **E.** Bias index (20-lap forward _rolling_ average) following muscimol injections. Same colors as in D

